# Collective Behavior in Medaka Fish Depends on Discrete Kinematic States of Swimming Behavior

**DOI:** 10.1101/2025.07.25.666863

**Authors:** Roy Harpaz, Pedro Piquet, Yasuko Isoe, Mark Fishman, Florian Engert

**Author notes:** Correspondence and requests for materials should be addressed to R.H.

## Abstract

Complex collective behaviors such as schooling are believed to emerge from simple, individual-level computations that translate incoming information from conspecifics into actions. Recently, it has been proposed that discrete behavioral modes, or internal states, may modulate these computations, affecting the resulting collective behaviors. Direct evidence for such hierarchical control remains limited due to challenges in inferring hidden perception-action computations and uncovering discrete behavioral modes from continuous behaviors.

To address this, we analyzed swimming behaviors of Medaka fish (*Oryzias latipes*) throughout development. At the group level, Medaka exhibit synchronized swimming formations that develop early, emerging around two weeks of age and stabilizing within one month. Unlike many teleost species that use burst-and-coast swim patterns, Medaka exhibit continuous tail and body undulations. We show that this continuous behavior can be segmented into three distinct kinematic states: acceleration, deceleration, and prolonged constant speed swimming. Using state-dependent computational models, we tested how Medaka translate social information from neighbors into actions across these kinematic states. The models revealed distinct computations governing social information processing and decision making in each state. Moreover, social responsiveness varied significantly between states—it was strongest during constant-speed epochs, intermediate during accelerations, and lowest during decelerations. Compared to similarly-sized zebrafish that employ burst-and-coast kinematics, Medaka exhibited greater diversity in state-dependent social interaction computations, ultimately resulting in stronger coordinated swimming.

These findings highlight discrete behavioral modes as key modulators of social interaction computations underlying collective behavior.

## Introduction

Complex collective behaviors such as schooling and flocking are critical for animal survival ^1^. The prevailing theory suggests that these collective behaviors are the emergent result of a set of decentralized, ‘simple’, local computations translating social information into action, which are considered static and even genetically hardwired ^2–4^. Previously, extensive theoretical ^2,3,5–8^ and experimental ^9–14^ studies have aimed to elucidate the nature of social interaction computations and the resulting collective behaviors. However, the exact nature of these computations and whether general principles apply across different species and taxa remain under debate ^13,15,16^.

Recent studies have suggested that the computations animals use to respond to social information are themselves modulated by internal and external factors. Environmental cues such as the presence of food or predators^17,18^, spatial confinement^19,20^, or changes in light intensity^21^ have been reported as modulators of social interactions. Furthermore, the experience of different social densities was recently reported to modulate the strength of responses to visual social information in young zebrafish, on the time scales of minutes to hours^22^. Such dynamic responses to external factors are hypothesized to help fish adapt effectively to changing environmental conditions ^22^.

Further, it has been suggested that animals frequently alternate, on the order of seconds, between discrete behavioral modes that modulate their social interaction computations^23–26^. However, empirical evidence supporting this hierarchical control is limited, primarily due to the challenge of simultaneously inferring hidden interindividual computations and discrete behavioral modes from continuous behavioral observations of multiple individuals.

One exception is the case of intermittent motion. Recent evidence indicates that different kinematic swimming modes influence interindividual interactions and emergent collective behaviors. For example, zebrafish larvae move in discrete bouts interspersed by times of inactivity ^27–29^. This naturally enables parsing behavior into discrete decision events. It is hypothesized that these animals integrate social (and all other types of information) primarily during the periods of inactivity^14,27^. Further, studies of adult zebrafish, that utilize burst-and-coast kinematics (but rarely show complete inactivity like larvae), demonstrated that social interactions can be similarly parsed into discrete ‘active’ and ‘passive’ processing modes, that correlate strongly with their acceleration and deceleration kinematics ^23,24^. Burst-and-coast kinematics was also found to be crucial in shaping the timing of responses to social information in young zebrafish ^25,26^. Similarly, intermittent stop-and-go motion was found to shape the collective marching of locusts ^30–32^. Lastly, various theoretical models have been developed to investigate collective behaviors in species that rely on intermittent motion ^11,33–35^.

However, intermittent motion is only one broad class of animal motion^36^. In fish, a common classification differentiates between intermittent (e.g. burst-and-coast) and continuous undulatory swimming, where different species tend to predominantly utilize one mode over the other ^37–41^. Intermittent swimming has been associated with reduced energetic expenditure ^38,42^ (but evidence is mixed ^41,43^), and enhanced sensory capabilities ^44,45^. Continuous swimming has been associated with energetic benefits at sustained swimming speeds ^43^ and improved oxygen availability ^46^.

Currently, less is known about how continuous motion patterns are related to discrete behavioral movement decisions, and how they affect collective group behaviors. Recent studies have successfully parsed continuous behaviors of individuals from various species (e.g., worms, flies and mice) into discrete behavioral states using their observable movement kinematics ^47–49^. Successfully parsing continuous motion into discrete behavioral modes during social interactions can facilitate the identification of alternative sets of social interaction computations underlying the resulting collective behaviors. However, such parsing and analysis have not yet been systematically applied to continuous-swimming fish, where continuous motion could strongly influence social interactions and collective behaviors.

To address this gap, we analyzed the individual and collective behavior of the continuous-swimming species, Medaka (*Oryzias latipes*), from early larval stages through adulthood ^40,50–52^. We show that Medaka swim in synchronised formations that emerge early ^53^, around two weeks of age, rapidly reaching stable adult levels by the juvenile age of one month. By analyzing individual speed profiles, we identified three discrete, non-overlapping swimming states: active acceleration, active deceleration, and a constant-speed state where fish maintain moderate speeds for extended periods. Using models that predict an individual’s swimming velocities based on the spatiotemporal kinematic features of its neighbors, we found that fish utilize distinct social interaction computations in each of the identified kinematic states. Moreover, prediction accuracy was notably higher during the constant speed state, indicating times of enhanced responsiveness to social information. Interestingly, compared to similarly sized fish species employing burst-and-coast kinematics, such as zebrafish, Medaka demonstrated greater diversity in state-dependent social interactions and stronger coordinated swimming. Our findings indicate that continuous undulatory swimmers utilize a diversity of social interaction computations which are strongly correlated with their kinematic swim states. Such diversity is hypothesized to contribute to the strong sociality of this fish species.

## Results

### Collective behavior in Medaka develops early and saturates rapidly

To study individual and collective behavior in continuous undulatory swimmers, we analyzed the swimming behavior of groups of the Japanese rice fish (*Oryzias latipes*), known as Medaka (Fig. 1a-b Movie 1). We tracked groups through development and measured their behavior at the ages of 10, 16, 24 and 34 days post fertilization (dpf), and at the adult stage (>60 dpf). We estimated each fish’s position, heading orientation and the midline of the body, and calculated the relative distances, azimuths and heading directions of its neighbors (Fig. 1b-inset, Methods).

**Figure 1.**
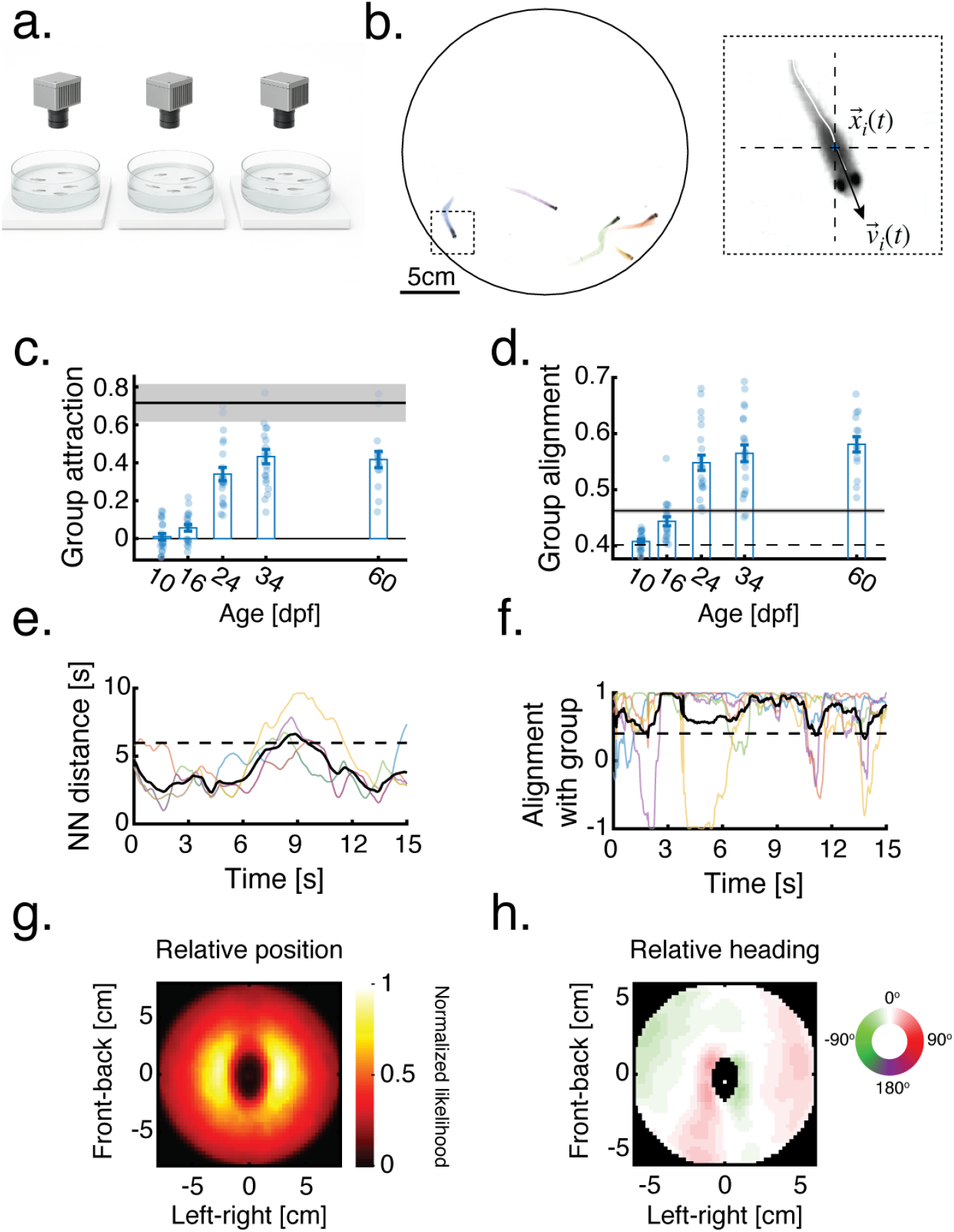
Social behavior in Medaka develops early and rapidly. **a**. Sketch of the experimental setups used for behavioral tracking of Medaka groups (Methods). **b**. A snapshot of a group of 5 adult Medaka (>60 dpf) swimming in a large arena (Diameter = 26cm). Different colors represent different fish in the group. **Inset**. Zoom in view of a single fish *i*, its center of mass position 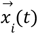, velocity vector 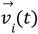, and its midline (bold white line). **c-d**. Average attraction (c.) and alignment (d.) values of Medaka groups (5 fish in a group) at different developmental stages. 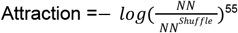, where *NN* is the average nearest neighbor distance, and *NN*^*Shuffle*^ is the expected value for shuffled groups (Methods).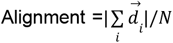, where 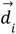 denotes the unit heading vector of fish *i* and *N* is the number of fish in the group. Dotted line is the expected value for shuffled groups. Bold horizontal black lines and shaded areas are the mean and SEM of attraction and alignment values for adult zebrafish groups (5 fish in a group, N=6 groups, Methods). **e**. Example traces of the NN distances for 5 adult Medaka (>60dpf) swimming in a group (colored lines), the average of the group (black line) and the expected value of shuffled groups (dotted black line). **f**. Example traces of the alignment of individual fish with the heading direction of the group (colored lines)(Methods), and the alignment of the group (black line). Same group as in e. **g**. Heatmap showing the likelihood of observing a neighboring fish at different spatial positions relative to a focal fish located at (0,0) pointing north. Groups of 5 fish (age >60 dpf). **h**. Average relative heading of all neighbors at different spatial locations with respect to a focal fish located at (0,0) pointing north. White colors represent no deviation or perfect alignment. Same groups as in g. In c. and d. dots represent average values of different groups, and error bars are SEM. Number of Medaka groups tested at different ages are *N*_10_ = 19, *N*_16_ = 20, *N*_24_ = 22, *N*_34_ = 22, *N*_60_ = 15.

We find that social behavior develops early in these fish, with signs of attraction emerging already at 16 dpf (8 days post hatching, Methods), and monotonically increasing as the fish mature (group attraction = 0.01±0.08, 0.06±0.08, 0.34±0.16, 0.43±0.18, 0.42±0.16 [mean±sd] for ages 10, 16, 24, 34, and >60 dpf)(Fig. 1c, e, Methods). Alignment with neighbors follows a similar developmental trajectory (group alignment = 0.41±0.02, 0.44±0.04, 0.55±0.06, 0.56± 0.07, 0.58±0.05)(Fig. 1d,f, Methods), and both properties are close to the adult values at the juvenile age of 34 dpf. Compared to adult zebrafish—a similarly sized teleost commonly studied for collective behaviors—Medaka groups exhibit more strongly aligned formations but reduced group cohesiveness (Fig. 1c–d), indicating a stronger tendency toward schooling rather than shoaling behaviors ^54^.

Analyzing group structure at the age of 60 dpf, when social behavior is fully developed, revealed the unique spatial organization of medaka groups: neighbors are more likely to be found directly to the side of a focal fish at intermediate distances (~2.5cm), but are less likely to be found directly in front or behind the focal fish (Fig. 1g). Alignment of heading direction is similarly high in areas corresponding to the most likely positions of neighbors, and also directly in front of the focal fish, emphasizing the anisotropic structure of the group. (Fig. 1h).

These results confirm that Medaka show distinct and robust social behaviors in a group setting, which are fully developed at the age of 60 dpf, allowing us to study the social interaction computations underlying collective behaviors at this age.

### Medaka display distinct kinematic swim states

We next analyzed the unique swim kinematics of adult Medaka (age 60 dpf). As reported before ^40^, Medaka swims with continuous tail and body undulations, as opposed to the punctuated burst-and-coast kinematics utilized by zebrafish for example (Fig. 2a, S1a-c, Movies 2-3). Importantly, we were able to naturally segment the speed profile of Medaka into three distinct kinematic states - acceleration and deceleration phases where fish modulate their swimming speed upwards or downwards, and a constant speed state in which fish maintain constant velocity magnitudes (Fig. 2b, Methods). To confirm that these three states are distinct, we demonstrate that speed profiles in each state are characterized by a unique family of functions. Accelerations are best described by a sigmoidal function (*R*^2^ = 0. 99 ± 0. 001 [mean±sd]), decelerations by a single exponential (*R*^2^ = 0. 96± 0. 002)^23^, and the constant speed epochs by an oscillating sine wave (*R*^2^ = 0. 74± 0. 03)(Fig. 2b). Each function family provided a uniquely good fit for only one state and failed to describe the other states (Fig. S1d). Mechanistically, accelerations were characterized by a range of time constants τ_*Acc*_= 0.044±0.013 (mean±sd) that were independent of the amplitude of the swim epoch (Fig. 2c). Decelerations were also characterized by a range of time constants τ_*Dec*_=0.39±0.16, with large amplitude epochs having shorter time constants (Spearman’s correlation = −0.43, p<0.0001, permutation test). During constant speed epochs, speed oscillated at 3.34±1.34 Hz (mean±sd), slower than the measured tail beat frequencies (Fig. S1a), with narrow amplitude variations around the mean speed (0.35±0.13 [mean±sd])(Fig. 2c, h). Higher oscillation frequencies tended to have lower amplitudes (Spearman’s correlation = −0.48, p<0.0001, permutation test).

**Figure 2.**
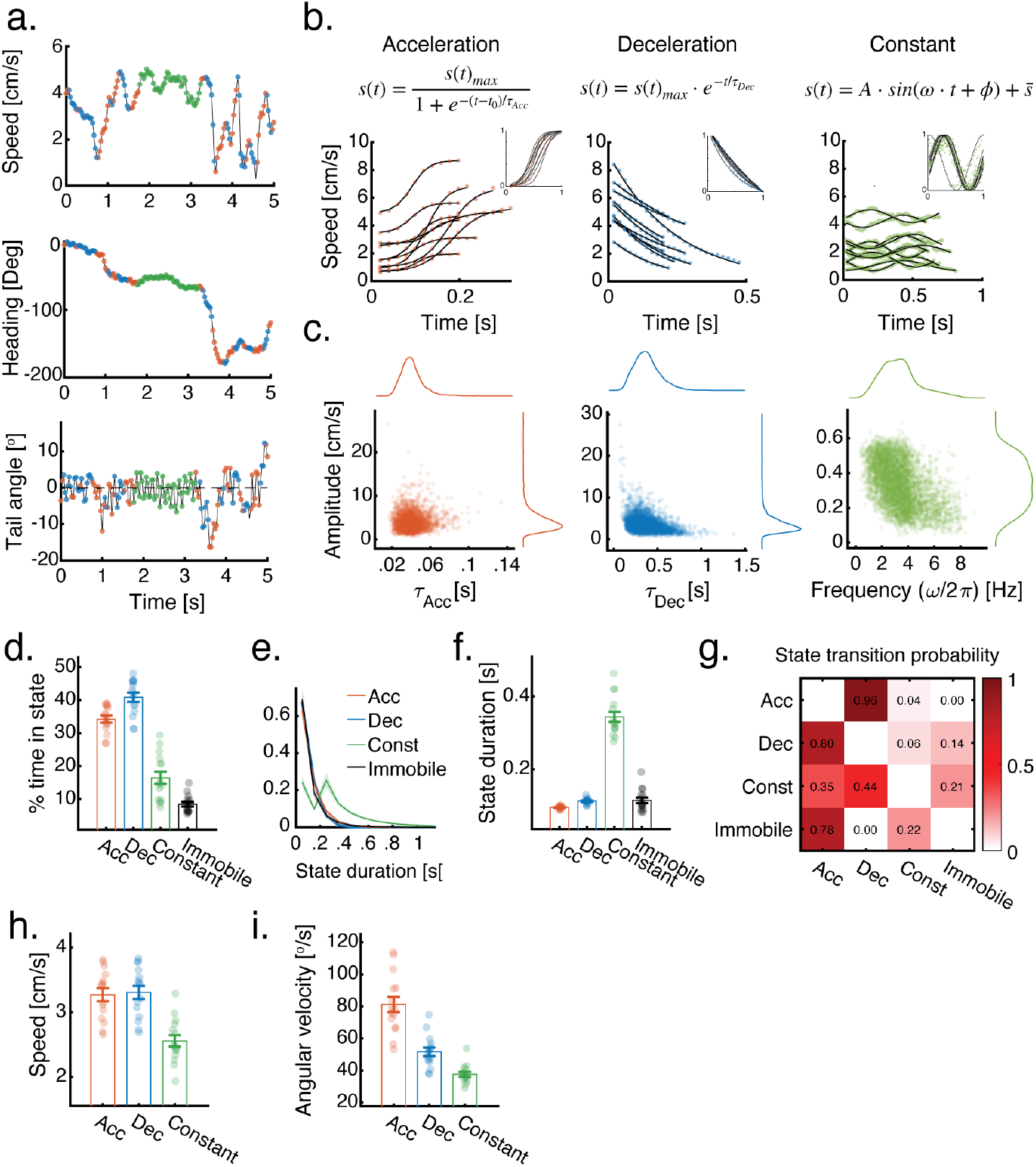
Medaka utilize distinct kinematic swim states. **a**. Examples of the speed, heading direction and tail angle deviation from the body axis of an adult Medaka swimming in a group of five. Red, blue and green colors represent acceleration, deceleration and constant speed epochs (Methods). **b**. Examples of speed traces showing acceleration, deceleration and constant speed epochs (colored dots) along with their best fits (black lines) when using sigmoidal, exponential and sine wave functions respectively. **Inset**. Same fits as shown in the main figures, but with time and speed magnitude normalized between 0-1. Normalized sine waves were also inverted to align the direction of their initial oscillation. **c**. Function parameters inferred from fitting segmented speed epochs (as in b.) plotted vs. the amplitude (|max-min|) of the epochs. Colored lines are marginal distributions. **d**. Average percent time spent in each of the segmented states for groups of adult Medaka. **e**. Distribution of state durations for each of the kinematic states. Bold lines are averages over fish, shaded areas are standard deviations. Bin width = 0.1s. **f**. Average duration of the segmented states (same groups as d). **g**. Transition matrix showing mean probability of transitions from a given state (rows) to each of the other states (columns). Probabilities are calculated over rows, averages are over groups. **h-i**. Speed (h.) and Angular velocity (i.) of adult Medaka in the three kinematic states. In d, f, h, and i, dots represent average values of different groups, and error bars are mean±SEM.

Next, we analyzed how these discrete states shape the swimming behavior of the fish. Adult Medaka spent 16.4±7.1% (mean±sd) of their swim time in the constant speed state, compared to 34.2±4.1% and 40.9±5.5% in the acceleration and deceleration states (Fig. 2d). The average duration of constant speed epochs, however, was ~3.6 and 3 times longer than acceleration and deceleration events respectively (Fig. 2e-f). In contrast, adult zebrafish, which are known to utilize ‘burst-and-coast’ kinematics, are rarely found in a similar constant speed state (<3%)(Fig. S1b-c, Movie 3). Analyzing the transitions between states, we find that Medaka frequently alternate between acceleration and deceleration states, transitioning less frequently into the constant speed state, which was sustained for longer durations (Fig. 2g, S1e).

Average swimming speed was similar across acceleration and deceleration events (Speed_Acc_=3.27±0.39, Speed_Dec_=3.31±0.39 cm/s [mean±sd])(Fig. 2h). Yet, speed was significantly slower in the constant speed state (Speed_Const_=2.55±0.34 cm/s), indicating that fish maintain slower speeds for longer durations. Turning was most strongly associated with accelerations and weakest during constant speed epochs (Turning_Acc_=81±18, Turning_dec_=52± 10,Turning_Const_=38±6 deg/s [mean±sd])(Fig. 2i, Methods). We did not observe any differences in distances and angles to the walls of the arena between the three states (Fig. S1f).

Together, these findings demonstrate that while Medaka utilize continuous body and tail undulations for swimming, they clearly employ at least three discrete kinematic states to regulate their swimming behaviors. Next, we assessed whether these kinematic states also influence the social interactions underlying collective behaviors.

### Distinct swim states imply varying levels of social responsiveness

Next, we study the relation between the distinct kinematic swim states of Medaka and their responses to social information from their neighbors. Based on the measured spatial structure of neighbor positions and relative headings (Fig. 1g-h), we opted to utilize a previously suggested family of models in which the change in velocity 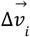 of fish *i* depends on the spatio-temporal properties of its neighbors ^23^. Specifically, we decompose the change in velocity of fish *i* at time *t* into two components (Fig. 3a):

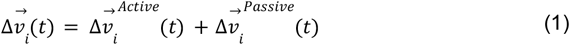

**Figure 3.**
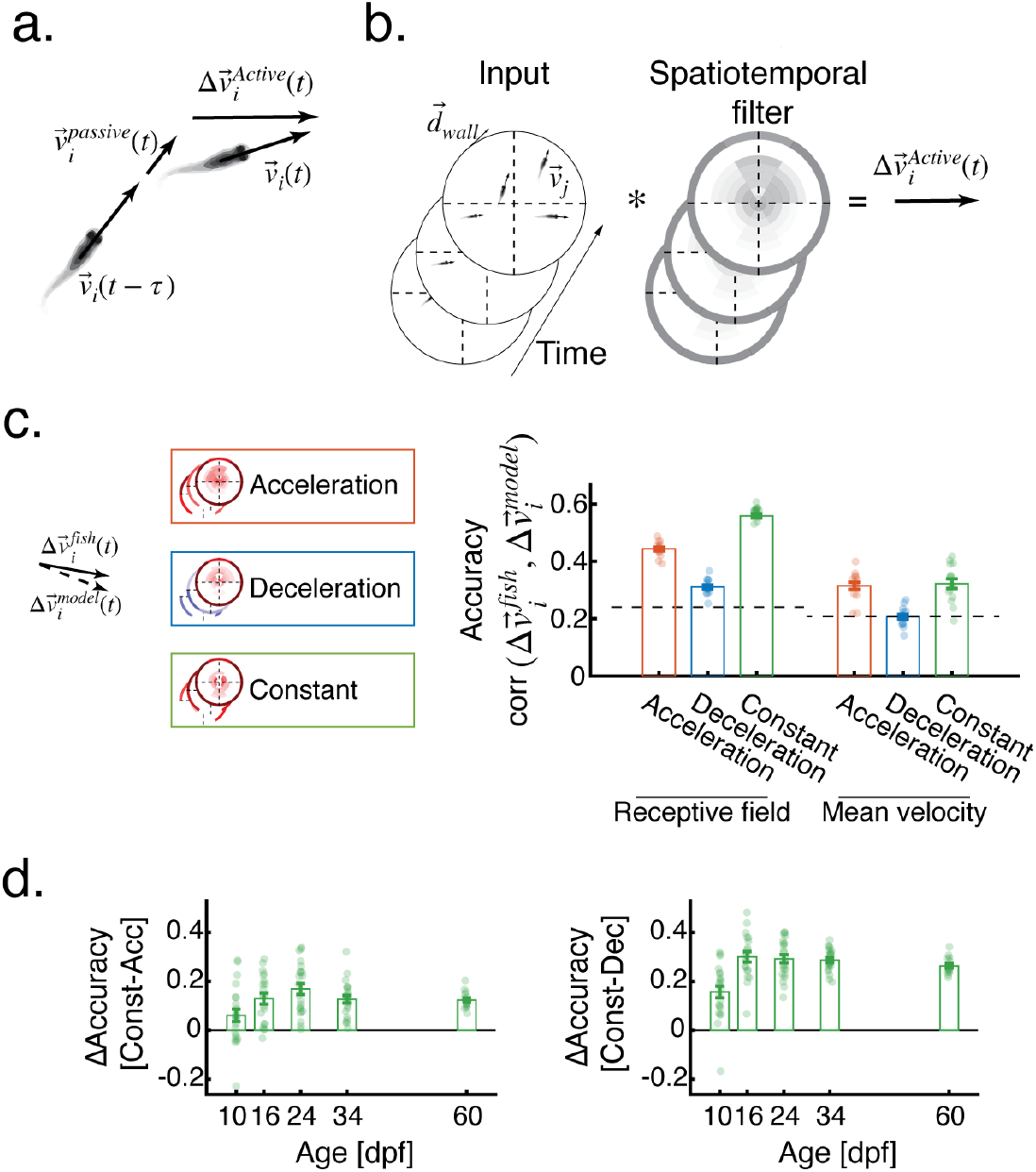
Distinct swim states imply varying levels of social responsiveness. **a**. The velocity fish *i*, can be decomposed into passive and active components (Eq. 1-2). **b**. The active motion component 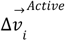 is modeled as the weighted spatiotemporal integration of neighboring fish velocities 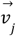 and the direction of the closest walls of the arena 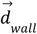 (eq. 3). Grayscale values represent different weights for different spatiotemporal bins; the outer ring represents weights for arena walls. **c**. Prediction accuracy measured as the correlation between the active motion component of the fish 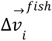 and the predicted active component by the models 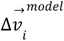. Models are learned separately for each state (see Fig. 4a) and accuracy is calculated using held out test data not used for training. ‘Receptive field’ is the full spatiotemporal model presented in b., and ‘Mean velocity’ is a simpler model that ignores the spatial positions of neighbors (Methods). Black dashed lines represent the accuracy of models fitted to shuffled data (Methods). **d**. Difference in prediction accuracy between the constant speed state and the acceleration (left) and the deceleration (right) states for different ages when using the full spatiotemporal models for prediction. In c and d, dots represent groups and error bars are mean±SEM.

The passive component represent velocity decay due to water friction, and is defined as:

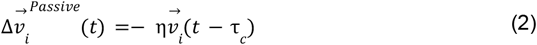

where η is the friction coefficient taken as the inverse of the measured time constant of the exponential fits of deceleration segments 1/⟨τ _*Dec*_ ⟩(Fig. 2b-c), and τ _*c*_ is a fixed decay timescale, estimated from the autocorrelation function of the speed time series (Fig. S2a, Methods).

The active component 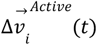 represents the response of the fish to social and sensory stimuli and is defined here as a spatiotemporal filter (or ‘receptive field’) of the velocities of neighboring fish and the direction of the closest walls of the arena:

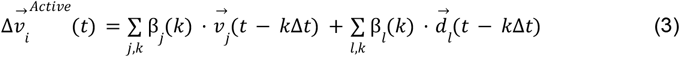

Where 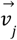 is the average velocity of neighbors in the spatiotemporal bin *j* at time *t* − *k*Δ*t*, and 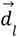 is the direction tangent to the wall closest to the fish. β _*j*_ (*k*) and β _*l*_ (*k*) are the weights assigned to each spatiotemporal bin for both social and sensory information respectively (Fig. 3b). In addition, we assume that a unique set of model weights β _*j*_ (*k*)^*state*^ and β _*l*_ (*k*)^*state*^ define each of the kinematic states of the fish - acceleration, deceleration and constant speed (Fig. 3c).

To test the ability of the models to predict fish active swimming responses, and to test for differences in social interactions between the states, we train a unique model for each group in each of the 3 kinematic states, and test how well the predicted motion vectors 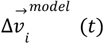 describe the actual changes in fish swim velocities 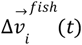 (Fig. 3c). While all states show above chance prediction accuracies (defined as the correlation between real and predicted velocities), we find clear differences between the state dependent models: Deceleration epochs showed the lowest prediction accuracies (0.31±0.02 [mean±sd], Pearson’s correlation coefficient), acceleration events were predicted with higher accuracies (0.44±0.02), and the constant speed epochs had the highest prediction accuracy of the three states (0.56±0.03). To estimate the importance of spatial information in the fish’s computations, we compared these accuracies to a baseline model where we predict the change in velocity of fish *i* as a simple average of neighbor velocities and wall directions regardless of their relative spatial positions (Eq. 5, Methods). This ‘Mean velocity’ model had markedly lower prediction accuracies compared to the full ‘Receptive field’ model (*C* _*acceleration*_ =0.31±0.05, *C* _*deceleration*_ =0.2±0.034, *C*_*const*_=0.31±0.06 [mean±sd], Pearson’s correlation coefficient), and no difference in accuracy between acceleration and constant speed epochs (Fig. 3c), emphasizing the importance of spatiotemporal information in the fish’s social interaction computations.

Importantly, segmenting fish swimming into only two states (acceleration and deceleration only, Methods) in the context of the full spatiotemporal model, ‘averages out’ the improvements in accuracy we observed in the constant speed state, resulting in prediction accuracies that are similar to those found for deceleration and acceleration events in the 3 state models (Fig S2b). Training a single model for the entire dataset, again results in intermediate accuracy values which are expected due to averaging (Fig. S2b), emphasizing the importance of the different social interaction states.

To confirm that prediction accuracies indeed depend on unique social interactions in the different kinematic states, we created a ‘shuffled identity’ dataset that retains the individual swim kinematics of the fish, yet randomizes available social information, by mixing fish trajectories from different groups (Methods). We find that models trained on this shuffled dataset, yield prediction accuracies that are at chance levels with almost no noticeable differences between the different kinematic states (Fig. S2c). This indicates that differences in the specific swim properties in each state are not enough to yield considerable difference in prediction accuracies, but rather depend on the unique social interactions at each kinematic state. As expected, fitting similar models to groups of adult zebrafish, which rarely display constant speed states, failed to result in an increase in prediction accuracy in the constant speed epochs, and showed lower overall model accuracies (Fig. S2d). Lastly, we found that the improved prediction accuracy in the constant speed state compared to the acceleration and deceleration states was consistent throughout development. Specifically, models fitted for individual groups, at different ages (Fig. S2e), showed an increase in accuracy in the constant speed state over the deceleration states (Δ*Accuracy*_*Const*−*Dec*_ = 0.16±0.11, 0.3±0.09, 0.29±0.07,0.29±0.05, 0.26±0.03 [mean±sd] for ages 10, 16, 24, 34, and 60 dpf respectively) and the acceleration states (Δ*Accuracy*_*Const*−*Acc*_ = 0.06±0.12, 0.13±0.1, 0.17±0.11,0.13±0.07, 0.12±0.04 [mean±sd]) from very early in development, and became stable around 24 dpf onwards (Fig. 3d).

Taken together, these results indicate that the three kinematic swim states are indicative of different social information processing states, with the continuous constant speed state showing the strongest social interaction tendencies.

### Fish utilize different spatiotemporal filters in each of the kinematic states

We next assessed the structure of the spatiotemporal filters (receptive fields) fish use in each of the kinematic states (Fig. 4a). In all states, the magnitude of the weights assigned to neighbor velocities was strongest at short temporal delays and gradually decreased as the temporal offset between neighbor velocities and focal fish responses increased (compare delays of −160 ms, −390 ms, and −630 ms in Fig. 4a). Spatially, the learned weights in the acceleration state extended up to 6 body lengths directly in front of the fish (Fig. 4a, top row, first column, 0° sector) but were weaker behind the focal fish (sectors 180°, 120°, −120°). By contrast, in the constant speed state, high-magnitude weights were tightly localized, reaching only about 2–3 body lengths in front of and to the sides of the focal fish, but extended farther behind it (Fig. 4a, bottom row, left column). Deceleration events had the weakest overall weight magnitudes, with a spatial structure similar to that of acceleration events (Fig. 4a, middle row, left column). These differences in learned weights, together with the fact that group structure did not markedly differ between states (Fig. S3a), indicate that fish differentially attend to their social surroundings in each kinematic state. Responses to arena walls were strong in all states, consistent with previous reports in other fish species ^23,56^.

**Figure 4.**
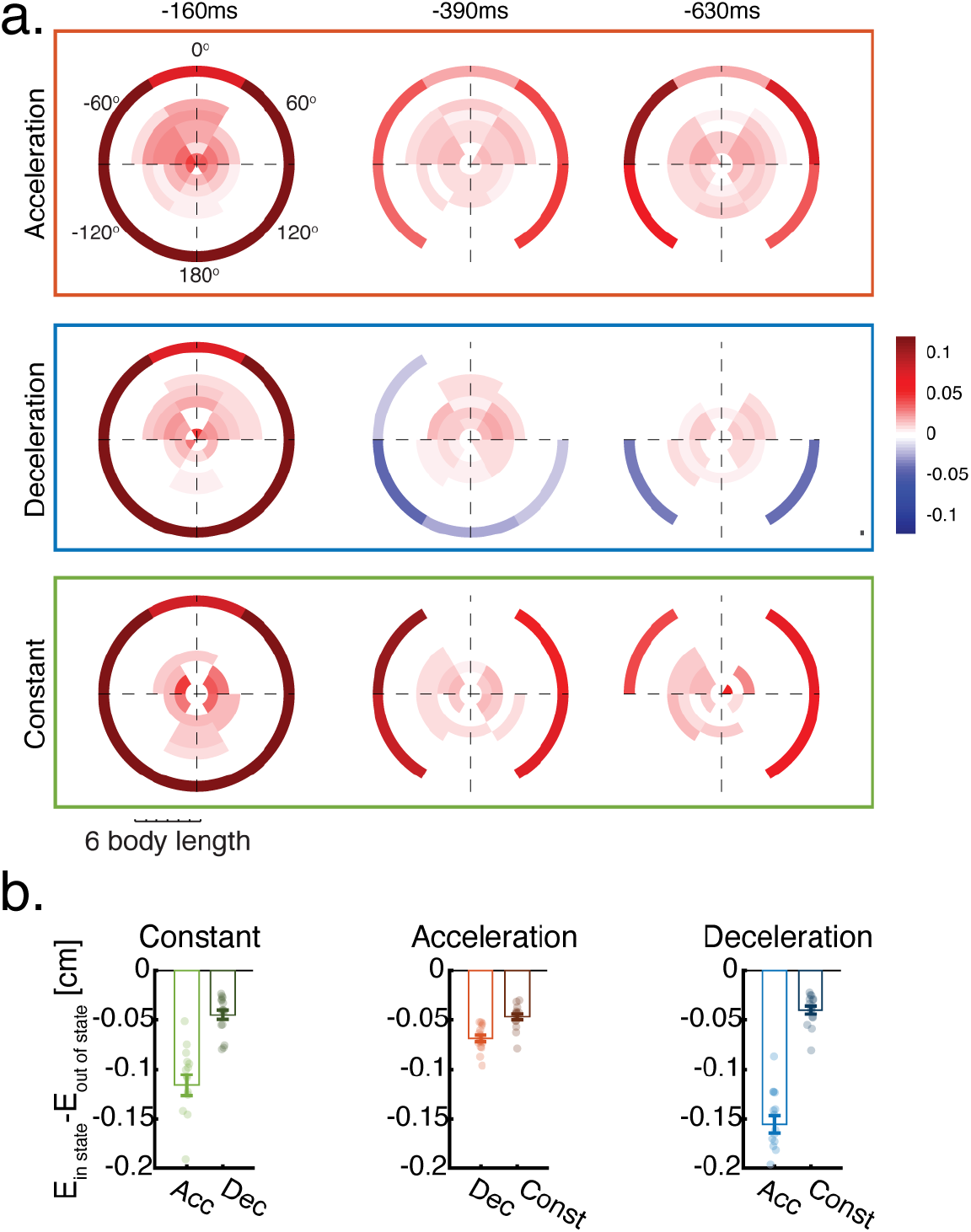
Fish utilize different spatiotemporal filters in each of the kinematic states. **a**. Learned spatiotemporal weights for the three kinematic states (rows). Each circular map represents the weights assigned to neighbor velocities at a specific distance and angle from a focal fish situated in the center of the map pointing north. Columns show maps at increasing Δ*t* offsets. The outer ring in each map represents the weights assigned to the wall of the arena at different angles. We only show weights that are significantly different from 0 (p<0.05, t-test, Methods). **b**. Difference in prediction error when using the model that was trained ‘in state’ (e.g. predicting held out velocities in the acceleration state using the model learned from acceleration data) and the predictions of the same velocities using models that were trained ‘Out of state’, i.e. on different kinematic states. Error is the Euclidean distance between the real velocity of fish *i* at time *t* and the predicted velocity by the models 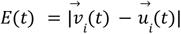. Average **‘**in state’ predictions are always better, regardless of the specific group or state analyzed, indicating the state specificity of learned spatiotemporal filters. dots represent groups and error bars are mean±SEM.

To confirm that fish indeed employ unique spatiotemporal filters in each kinematic state and to estimate their effect on behavior, we compared the magnitude of prediction error, defined as 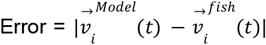, when using each of the three learned models shown in Fig. 4a to predict changes in velocity 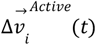 in each of the states. We show that prediction error is consistently and significantly lower when the model used for prediction matches the kinematic state of the target velocity vector, as evaluated on held-out data across all states (Fig. 4b). For example, predicting target velocities in the constant speed epochs using models learned from acceleration or deceleration epochs, resulted in a 0.11 ± 0.04 cm and 0.05 ± 0.02 cm (mean ± sd) increase in error, respectively, compared to the predictions of models trained on constant-speed data (Fig. 4b, left). This pattern is consistent across all kinematic states and experimental groups (Fig. 4b).

In summary, the distinct models learned for each state confirm that fish process and respond to spatiotemporal social information in a state-dependent manner.

## Discussion

In the current study, we demonstrated that the continuous undulatory swimmer, Medaka, exhibits robust collective behaviors that develop early—around two weeks of age—and rapidly stabilize, reaching adult-level values by the juvenile age of one month. These findings establish Medaka as a valuable model species for studying the development and mechanisms of collective social behaviors. Furthermore, we show that the unique kinematic profile of continuous swimming can be segmented into discrete states, each strongly associated with distinct social interaction computations. This diversity of social interaction patterns enhances the animals’ responsiveness to social cues, ultimately shaping their collective behavior.

The discrete, non-overlapping kinematic states we detected provide strong predictions about the levels of social responsiveness in Medaka. While accelerations are expected to be strongly related to executed behavioral decisions, and decelerations to reduced responsiveness to neighbors ^11,14,23,34^, the marked increase in responsiveness to social information in the constant speed state has not been described previously. We hypothesize that maintaining slow, constant swim speeds enhances responsiveness to social information, possibly due to enhanced sensorimotor processing during prolonged steady swimming. Future studies could test whether such improved responsiveness is specific to social information or represents a general enhancement of sensory processing, including responsiveness to non-social stimuli ^57–60^.

Furthermore, it is likely that additional behavioral states, beyond the three states identified in this study, influence the social behaviors of this species^27,61^. For example, fish may utilize different behavioral rules, or modify the usage of these kinematic states during specific contexts such as when foraging for food ^18,27,40^ or in the presence of predators ^17,62^. Moreover, current or previous social densities and the social experiences of the animals might modulate behavioral states of individuals either in a discrete or continuous manner ^22,63^. Future research can now systematically explore how these various factors shape Medaka collective behaviors.

The ontogeny of collective social behavior in Medaka can be directly compared with previous findings in zebrafish, another freshwater fish species of a similar size ^10,14,64,65^. Zebrafish larvae exhibit predominantly repulsive interactions at a young age of ~7dpf, developing attraction tendencies later, between 14 and 21 dpf ^14,22,55^. In contrast, Medaka larvae show no signs of purely repulsive tendencies at early ages, with robust social attraction developing already at 16 dpf (Fig. 1c). Moreover, the characteristic ‘burst-and-coast’ kinematics of adult zebrafish have previously been segmented into ‘active’ and ‘passive’ social interaction states^23^. In comparison, Medaka’s continuous swimming kinematics exhibit a greater diversity of social interaction states, all classified as ‘active’, even decelerations. We hypothesize that these differences underlie distinct collective behaviors between species, namely the stronger schooling observed in Medaka compared to the predominantly shoaling behavior in zebrafish. Importantly, since behavior in these species diverges already at the young larval stages, when brain-wide calcium imaging and optogenetics circuit dissection are widely available^66–69^, our findings provide an opportunity to study differences and commonalities in the underlying circuit mechanisms^70^.

To analyze the social interaction computations underlying collective behaviors in Medaka, we adapted a family of ‘receptive field’ models previously validated in adult zebrafish^23^. Despite considerable behavioral differences at both individual and collective levels, we found that this modeling framework effectively captures Medaka social responses, emphasizing the general applicability of these models in studying collective behaviors. The fitted models allowed us to characterize how fish utilize spatiotemporal information from neighbors, revealing distinct receptive fields for each kinematic state. These insights can inform future theoretical studies investigating how diverse social interaction states shape collective behaviors in continuous undulatory swimmers, as was previously done for fish utilizing burst-and-coast swimming ^11,34,71,72^.

Recent technological advances have positioned Medaka as a promising model system for linking neural circuit mechanisms and emergent adaptive behaviors ^70,73–75^. Our results pave the way for studying such mechanisms in the context of social and collective behaviors in this important model system.

## Acknowledgments

We thank all members of the Engert and Fishman labs for their support and advice throughout this project. We particularly thank Kumaresh Krishnan and Cristina Santoriello for their valuable assistance in establishing the behavioral setups, and Emily Chan for her help with Medaka breeding. We also thank Prof. Elad Schneidman for valuable discussions and insights. Finally, we thank Ed Soucy, Brett Graham, and Yuwei Li from the Neuroengineering Core Facility at Harvard’s Center for Brain Science for their technical support. Roy Harpaz received funding from Harvard Minds Brain and Behavior initiative. Florian Engert received funding from the National Institutes of Health (U19NS104653 and 1R01NS124017-01), and the Simons Foundation (SCGB 542973 and NC-GB-CULM-00003241-02).

## Author Contributions

RH, PP, YI, MCF and FE designed research, RH, PP and YI performed experiments, RH, PP and FE analyzed the data, RH, PP, YI, MCF and FE wrote the paper

## Supplementary figures

**Figure 1.**
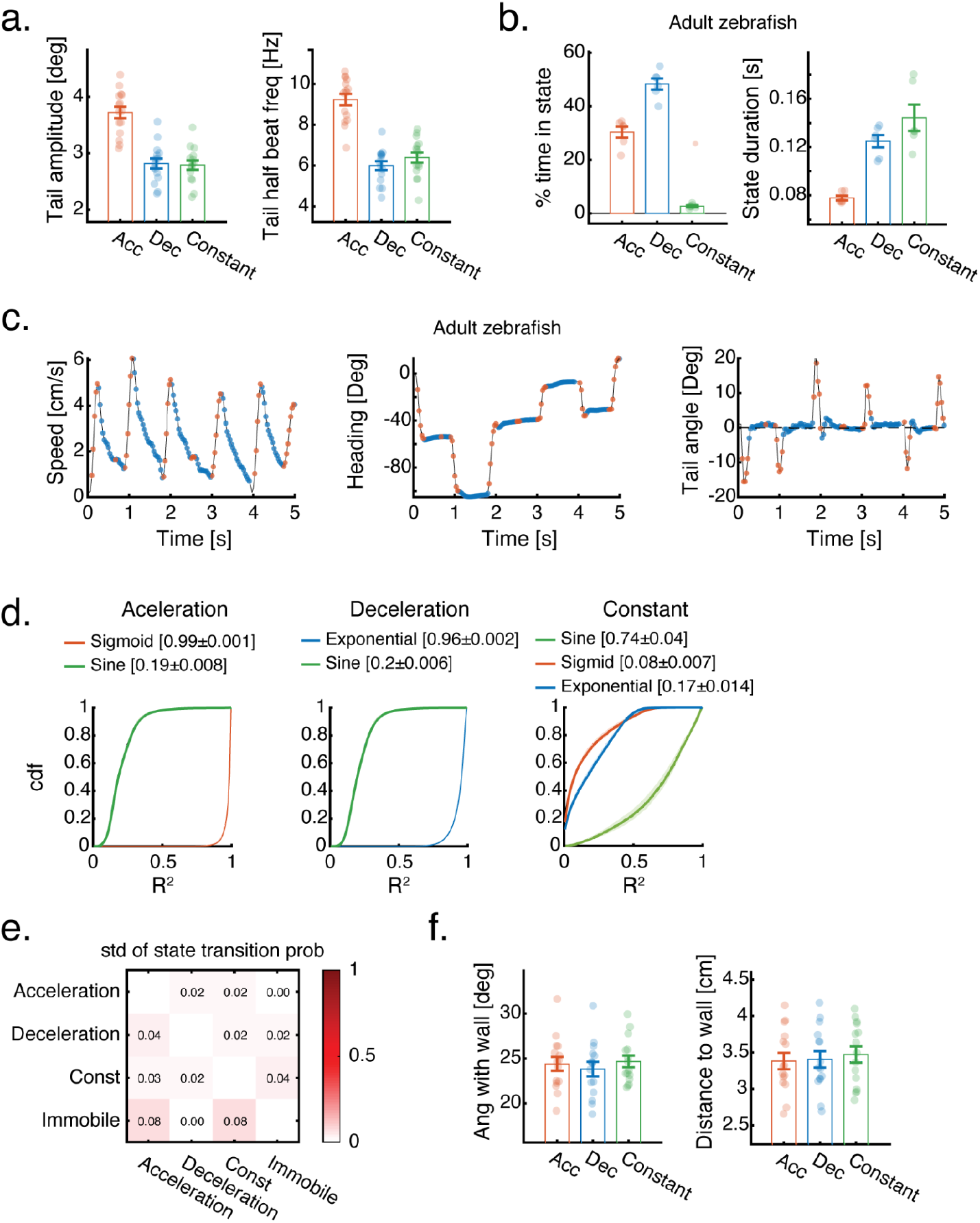
Distinct behavioral swim states of Medaka and zebrafish. **a**. Mean tail amplitude (left) and half beat frequency (right) of adult Medaka for the three kinematic states (Methods). **b**. Mean percent time spent in each of the kinematic states (left) and mean duration of the detected states (right) of adult zebrafish (Methods). **c**. Examples of the speed (left), heading angle (middle) and tail angle deviation from the body axis (right) of an adult zebrafish swimming in a group of 5 fish. Red and blue colors represent acceleration and deceleration events (constant speed epochs were not detected in this example). **d**. Cumulative density functions of goodness of fit values (R^2^) for acceleration (left), deceleration (middle) and constant speed epochs (right). Different colors represent the R^2^ values obtained by fitting different functions to epochs from a specific state. Bold lines and shaded areas are mean±SEM of the calculated cdf’s of different groups. Values in brackets are mean±sd. **e**. Standard deviations of the mean state-transition probabilities shown in Fig. 2g. **f**. Mean angle between fish heading direction and the direction tangent to the closest wall (left) and mean distance to the closest wall (right) for the three kinematic states of adult Medaka. In a, b and f, dots represent groups and error bars are mean±SEM.

**Figure 2.**
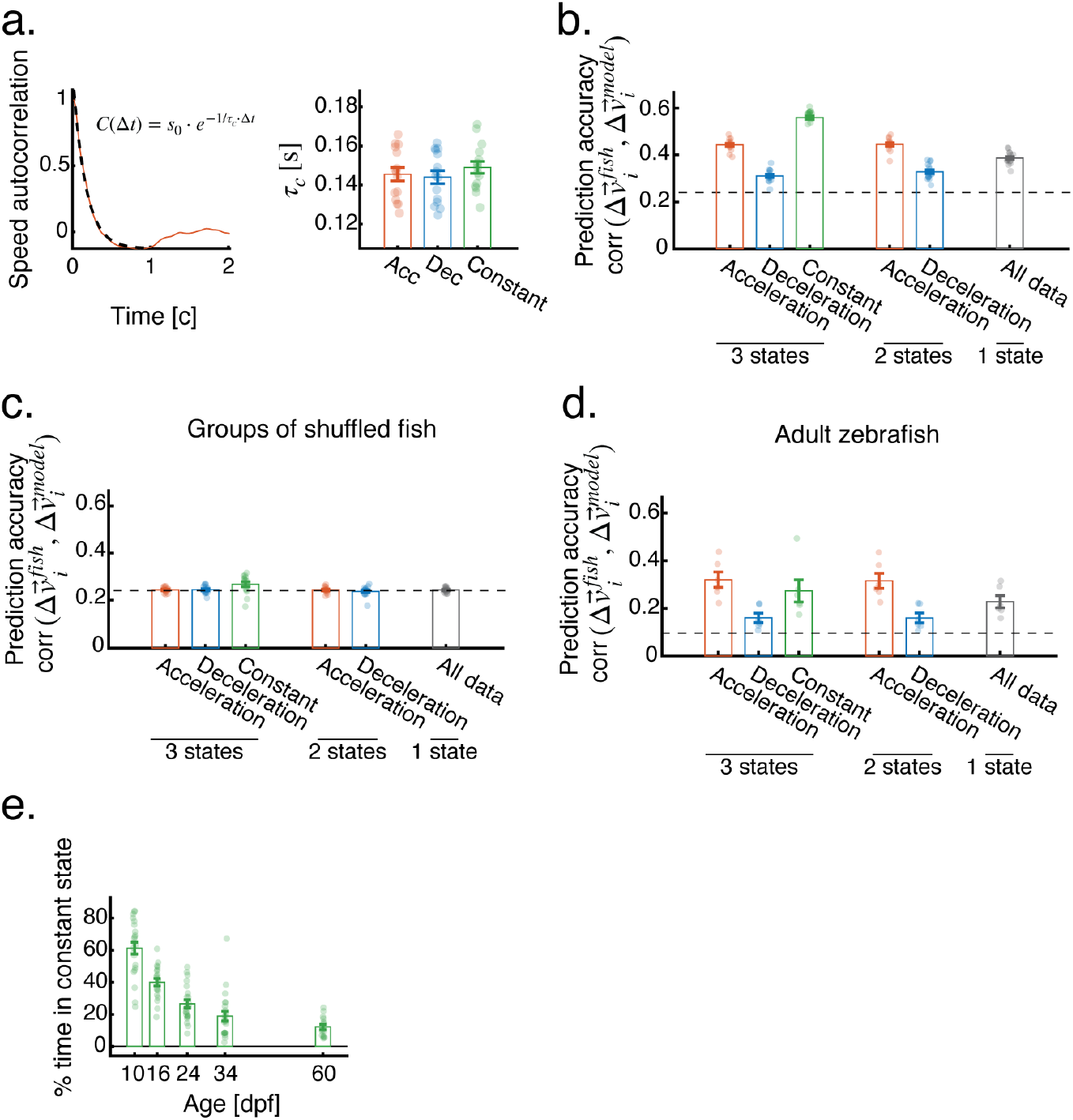
Distinct swim states imply varying levels of social interaction. **a. Left:** Example autocorrelation function (red) and the corresponding function fit of the form 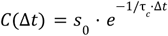 (black dashed line) calculated for acceleration events of an adult Medaka swimming in a group of 5. **Right:** Decay time constant τ _*c*_ of the auto-correlation function fits for each of the kinematic states. **b**. Prediction accuracy of the spatiotemporal models (eq. 3), similar to Fig. 3c, but for different segmentations of the data - 3 states, 2 states (no constant speed) or a single state. **c**. Prediction accuracy for spatiotemporal models fitted to surrogate data created by mixing fish from different groups (Methods). All analysis on surrogate data was done in the same manner as for real fish data (compare to Fig. 3c, S2b). The dashed black line represents prediction accuracy when models are fitted to data shuffled over groups and time (Methods). **d**. Prediction accuracy (similar to Fig. 3c) but for adult zebrafish groups. Note the lack of improvement in prediction accuracy for the constant speed state for fish performing burst-and-coast swimming. **e**. Percent of time spent in the constant speed state for Medaka groups at different ages. In all panels, dots represent groups and error bars are mean±SEM.

**Figure 3.**
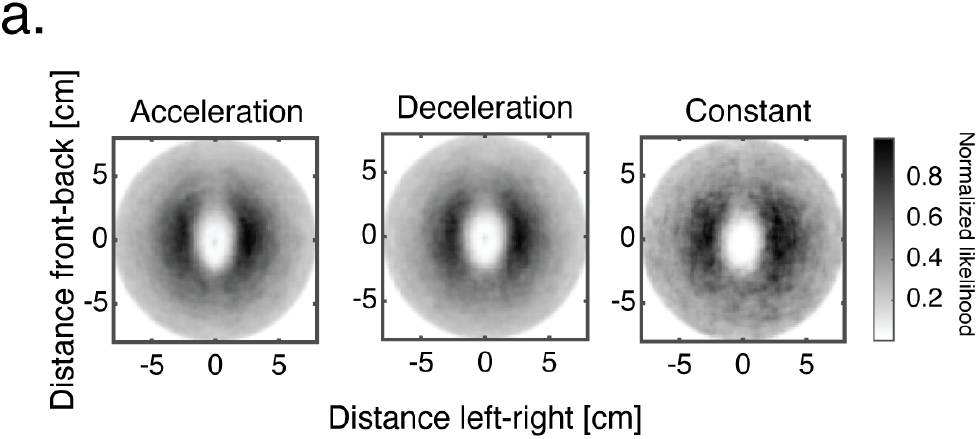
Spatial structure of groups is similar between the different kinematic states. **a**. Distribution of nearest neighbor positions with respect to a focal fish, situated at [0,0] pointing north.

## Methods

### Fish Husbandry

- **Medaka**. All fish used in the experiments were obtained from crosses of adult wild-type Medaka (drR). Larvae were raised in low densities of approximately 10 - 20 fish in medium petri dishes (D = 9 cm). Dishes were filled with filtered fish facility water and were kept at 28 °C, on a 14–10 h light-dark cycle. After hatching (8 dpf), fish were fed paramecia once a day. On day 2 after hatching, fish that were not tested in behavioral experiments, were returned to the fish facility where they were raised in 2 L tanks filled with 1.5′′ nursery water (2.5 ppt), with ~15 fish in each tank and no water flow. On days 10–12 water flow was turned on and fish were fed artemia 3 times a day until they were tested at subsequent ages.
- **zebrafish**. Experimental fish were obtained by crossing adult wild-type zebrafish (AB). Larvae were raised in conditions similar to Medaka. Fish were fed from 5 dpf onwards.

For both adult zebrafish and adult Medaka, male-to-female ratios were maintained at approximately 1:1 and individuals were randomly assigned to experimental groups.

All experiments followed institution IACUC protocols as determined by the Harvard University Faculty of Arts and Sciences standing committee on the use of animals in research and teaching.

### Behavioral experiments

- **juvenile and adult fish**. Juvenile Medaka (34 dpf), adult Medaka (>60 dpf), and adult zebrafish (>75 dpf) were transferred from their holding tanks to 3 arenas comprised of an opaque Polypropylene plastic pan (McMaster-Carr)(D = 26cm) filled with ~ 2-3 cm of water (depending on the size and age of the fish). Experimental arenas were filmed from above using an overhead Bassler acA2040-90umNIR camera equipped with Cinegon 1.9/10(0901) lens. Each arena was lit from below using a specialized infrared 24×24 inch IR light panel (850nm, Knema) and from above by indirect light coming from 4 32 W fluorescent lights. Recorded images were collected at ~50fps and tracked offline using software written in house using Matlab (see ^23^ for details).
- **Larval fish**. All behavioral experiments involving larval fish (10-24 dpf) followed the experimental protocols described previously ^14^. Briefly, fish were transferred from their holding tanks to custom-designed experimental arenas of sizes d = 6.5, 9.2, 12.6 cm (for 10, 16 and 24 dpf fish respectively), filled with filtered fish facility water up to a height of ~0.8 cm. Experimental arenas were made inhouse from sandblasted 1/ 16” clear PETG plastic and had a flat bottom and curved walls (half a sphere of radius 0.5 cm) to encourage fish to swim away from the walls. Every experimental arena was filmed using an overhead camera (Grasshopper3-NIR, FLIR System, Zoom 7000, 18–108 mm lens, Navitar) and a long pass filter (R72, Hoya). Arenas were lit from below using 4 infrared LED panels (940 nm panels, Cop Security) and from above by indirect light coming from 4 32 W fluorescent lights. Recorded images were immediately background subtracted and thresholded to segment fish bodies. The segmented binary images were saved for subsequent offline tracking and further processing. All acquisition and online segmentation were performed using custom-designed software written in Matlab ^14^.

Each group was imaged for 20 minutes, following 5–10 minutes of acclimation to the arena. Groups were eliminated from subsequent analysis in the case that one or more of the fish were immobile for more than 60% of the experiment ^14^. Only 3 groups of 10dpf larvae were eliminated from the experiment. Choosing a more stringent, or a less stringent criteria for elimination did not change the qualitative nature of the results.

#### Tracking

Segmented fish images were subsequently analyzed offline to extract the center of mass position 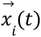 of each animal *i* at time *t*. Subsequent positions were then concatenated to obtain continuous individual fish trajectories^14,23^. Fish images were additionally analyzed to obtain the body orientation 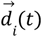 of the animals using second-order image moments, and to extract fish midline for tracking of body and tail undulations (see below).

#### Midline extraction and tail tracking

Binary fish images were analyzed to extract the midline of the body from head to tail. Briefly, we ‘skeletonize’ the binary image to get a single pixel width skeleton of the image. We detect branch points and endpoints along the skeleton, and choose the two endpoints that correspond to the edges of the image. We then used geodesic distances inside the skeleton to compute a shortest path connecting those endpoints. The resulting path was then smoothed and resampled to obtain the final midline of the fish (Fig. 1b, Movies 2-3).

#### Individual and group level swim properties

Fish velocity was calculated from the center of mass positions of the fish such that 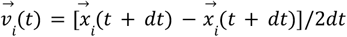, where dt is taken as one frame (~1/50 s). Speed is then defined as the magnitude of the velocity 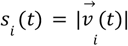. We measure group attraction as *Attraction*(*t*) = − *log*(*NN*(*t*)/*NN* ^*Shuffled*^), where *NN*(*t*) is the average nearest neighbor distance of the fish in a group and *NN*^*Shuffled*^ is the average nearest neighbor distance obtained from shuffling the identity and time signature of fish across groups (i.e. all fish in the shuffled groups are taken from different real groups)(Fig. 1c). Group alignment is defined as 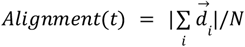, where 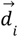 is the unit direction vector of fish *i*, and N is the number of fish in the group. Alignment is therefore bounded between 0 (if all fish point maximally away from each other, and 1, if all fish point in the same direction). For reference we also calculate the average alignment of shuffled fish groups (Fig. 1d). The alignment of a fish *i* with the direction of the group (Fig. 1f) is defined as the projection of 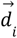 onto 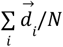, which is the average direction of the group, and this value is bounded between −1 and 1.

#### Neighbor Maps

Density maps (Fig. 1g) were obtained by calculating a 2d histogram of relative positions of all neighbors with respect to a focal individual situated at (0,0) pointing north. Maps were smoothed using a 2d rotationally symmetric gaussian filter with σ^2^ = 0.15cm. Scale Bar represents percent of the most visited bin. Angular deviation maps were calculated by collecting the direction of motion relative to a focal fish situated at (0,0) pointing north, of all neighbors that occupy a specific spatial bin around a focal fish. We then calculate an average direction vector for that bin and the angular deviation of that vector from north. Histograms were smoothed using a similar gaussian filter. For both maps, we discarded all data points in which fish were at a distance <0.3 Arena radii to the walls of the arena (3.9 cm for adult Medaka) to reduce the effects of arena boundaries on the structure of the maps.

#### Segmenting fish speed profiles into discrete states

We used the speed profile *s* _*i*_ (*t*) of each fish to segment fish swimming into acceleration, deceleration and constant speed states. We define a ‘constant speed’ epoch, as the longest segment we can detect that has similar start and end speeds, and low speed variability within the segment. We used the following algorithm: start from time *t* - and look for the farthest point *t* + *t* _*c*_ such that |*S*(*t*) − *S*(*t* _*c*_)| < δ_1_ and *variance*[*S*(*t*‥ *t*_*c*_)] < δ_2_, where δ_1_ = 0. 1*cm*/*s* and δ_1_ = 0. 2*cm*/*s*^2^. Segments had a minimal length of 150ms. Accelerations were then taken as all times that are not detected as constant speed epochs and had a positive acceleration sign, while deceleration epochs had a negative acceleration sign. Very low speeds (<0.6cm/s) were unassigned and were excluded from further analysis. Low speeds accounted for ~8% of all data.

For comparison, we also segmented fish speed profiles to only two states - accelerations and decelerations (Fig. S2b-d) according to the sign of the acceleration profile, similarly removing Very low speeds from further analysis.

#### Function fitting to kinematic states

Every identified speed epoch 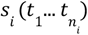 was assigned one of 3 labels - ‘acceleration’, ‘deceleration’ or ‘constant speed’ according to the algorithms described above. We then use the following function families to describe these epochs:

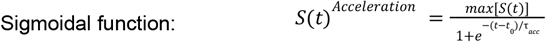

where *max*[*S*(*t*)] is the maximum speed in that epoch, *t*_0_ is the midpoint of the sigmoidal function and τ_*acc*_ is the time constant of acceleration. We optimize *t*_0_, and τ _*acc*_ for each acceleration epoch.

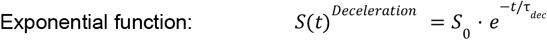

where *S*_0_ is the starting speed and τ_*dec*_ is the decay time constant. We optimize *S*_0_ and τ_*dec*_ for each deceleration epoch.

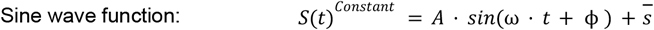

Where *A* is the amplitude, ω is the angular velocity, ϕ is a phase shift and 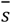 is the average speed of the epoch. We optimize *A*, ω and ϕ for each constant speed epoch.

For comparison, we also attempted to fit sine wave functions to acceleration and deceleration epochs, and sigmoidal and exponential functions to the constant speed epochs. These fits gave very low accuracy values showing that each family is uniquely apt at describing one state, but not the others.

#### ‘Receptive field’ models to predict individual velocity changes

We predict the active velocity changes of fish *i* at time 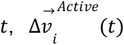, from the velocities of its neighbors in the recent past 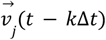 and their spatial configuration, as described in Eq. 3. To test for varying spatiotemporal effects in the different kinematic states, we fit a separate model for each of the kinematic states for every group tested. Model parameter values, or β weights, were learned using least squares Lasso regression ^76^, by minimizing the following cost function:

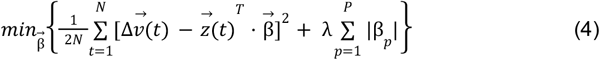

Where 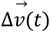 is the change in velocity of the fish, 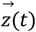 are neighbor velocities in the spatiotemporal bins around the focal fish, 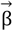 is the set of weights to be optimized, and N is the number of training observations. The second term in the parenthesis, penalizes the cost function for having many non-zeros parameters in the model, with hyper parameter λ controlling the magnitude of the penalty. Optimizing model parameters on held out test data usually results in a sparser model, setting non contributing weights to 0. We standardized all predictors 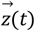 and target vectors 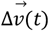 within each kinematic state and inside every cross-validation training fold to zero mean and unit variance before learning model parameters 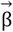, to ensure that differences in the scales or magnitude of the observations between the different states (Fig. 2h-i), will not bias model fit accuracies or the scale of the learned weights.

#### ‘Mean velocity’ model to predict individual velocity changes

To establish a baseline for model prediction accuracies, and to assess the unique contribution of spatiotemporal features in describing fish movement decisions, we also trained a simpler model that ignores the spatial properties of neighbors and only utilized their velocity information to predict the change in velocity of a focal fish 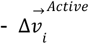 :

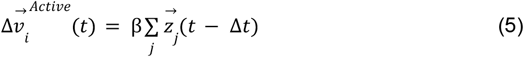

where 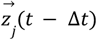 is the velocities of all neighboring fish and the direction of the closest wall 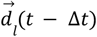 at time *t* − Δ*t*.

#### Surrogate data of shuffled groups

To confirm that prediction accuracies indeed depend on unique social interaction computations in the different kinematic states, and not on inherent difference of the observations between the states, we created a ‘shuffled identity’ dataset that retains the individual swim kinematics of the fish, yet randomizes available social information, by mixing fish trajectories from different groups. We then fit models to these dataset in the same way we fit data from real groups (Fig. S2c). Prediction accuracies from this surrogate dataset are similar to the ones obtained when we scramble the temporal contingency of observations and predictors and the spatial organization of neighbors velocities within groups (dashed black lines in Fig. 3c and S2b-c).

#### Model parameters

The following parameters were used in training state dependent spatiotemporal models (Eq. 3), according to values reported previously^23^.

| Parameter name | Description | Values used |
| --- | --- | --- |
| Distance bins | # of rings in each spatial filter | 6 bins, 0-6 body length at 1 body length increments |
| Angular bins | # of sectors in each spatial filter | 6 bins, covering 60 degrees each. |
| Time delays | # of spatial filters measured at different $\Delta t$ times into the past. | $\Delta t \approx -160\text{ms}$ (8 frames), $-390\text{ms}$ (20 frames), $-630\text{ms}$ (32 frames).<br>For the ‘simple average’ model we use $\Delta t \approx -160\text{ms}$ only. |

#### Weight map plots

β weight maps (Fig. 4a) show the average weights per spatiotemporal bin over all fitted groups and states. We perform a significance test for every spatiotemporal bin, using a one sample t-test, and only plot β weights with p-value < 0.05. Since our weight estimation procedure is already relatively conservative (weights are learned using a Lasso regularized cross-validation procedure for each group, which tends to shrink parameters towards 0), we do not perform additional post-hoc corrections.

#### Sample sizes, trial numbers and power estimation

Sample sizes were chosen based on previous experiments studying groups of larvae, juvenile and adult zebrafish for reference ^14,23,55^. These group sizes were expected to allow for accurate estimation of individual and group level properties and allow comparison between experimental conditions with sufficient statistical power.

#### Statistical testing

We used *R*^2^ values to assess goodness of fit measure when fitting functions to kinematic epochs (Fig. S1d). We used Spearman’s correlation coefficient to assess the monotonic relationship between the parameters of function fits and the amplitude of the speed profile (Fig. 2e), and estimated their statistical significance using a permutation test with 10000 shuffled repetitions (therefore the limit of p-value resolution is 1/10000). Finally we used a one sample t-test to detect estimated model weights that are statistically different from 0. Before performing a specific statistical procedure we made sure all model assumptions are fulfilled.

## Data availability

All data used in this study are publicly available at: https://dataverse.harvard.edu/dataset.xhtml?persistentId=doi:10.7910/DVN/QTNPJL.

## Notes

https://dataverse.harvard.edu/dataset.xhtml?persistentId=doi:10.7910/DVN/QTNPJL

### Competing Interest Statement

The authors have declared no competing interest.

